# Therapy Development for Microvillus Inclusion Disease using Patient-derived Enteroids

**DOI:** 10.1101/2023.01.28.526036

**Authors:** Meri Kalashyan, Krishnan Raghunathan, Haley Oller, Marie-Bayer Theres, Lissette Jimenez, Joseph T. Roland, Elena Kolobova, Susan J Hagen, Jeffrey D. Goldsmith, Mitchell D. Shub, James R. Goldenring, Izumi Kaji, Jay R. Thiagarajah

## Abstract

Microvillus Inclusion Disease (MVID), caused by loss-of-function mutations in the motor protein Myosin Vb (MYO5B), is a severe infantile disease characterized by diarrhea, malabsorption, and acid-base instability, requiring intensive parenteral support for nutritional and fluid management. Human patient-derived enteroids represent a model for investigation of monogenic epithelial disorders but are a rare resource from MVID patients. We developed human enteroids with different loss-of function MYO5B variants and showed that they recapitulated the structural changes found in native MVID enterocytes. Multiplex Immunofluorescence imaging of patient duodenal tissues revealed patient-specific changes in localization of brush border transporters. Functional analysis of electrolyte transport revealed profound loss of Na^+^/H^+^ exchange (NHE) activity in MVID patient enteroids with near-normal chloride secretion. The chloride channel-blocking anti-diarrheal drug, Crofelemer, dose-dependently inhibited agonist-mediated fluid secretion. MVID enteroids exhibited altered differentiation and maturation versus healthy enteroids. Inhibition of Notch signaling with the γ-secretase inhibitor, DAPT, recovered apical brush border structure and functional Na^+^/H^+^ exchange activity in MVID enteroids. Transcriptomic analysis revealed potential pathways involved in the rescue of MVID cells including serum- and glucocorticoid-induced protein kinase 2 (SGK2), and NHE regulatory factor 3 (NHERF3). These results demonstrate the utility of patient-derived enteroids for developing therapeutic approaches to MVID.

**Conflict-of-interest statement:** The authors have declared that no conflict of interest exists.

## INTRODUCTION

Microvillus Inclusion Disease (MVID) is a rare congenital diarrhea and enteropathy (CoDE) caused by loss-of-function mutations in the motor protein Myosin Vb (MYO5B) (1–5). In general, patients with severe loss of MYO5B function have profound diarrhea, intestinal fluid loss and nutrient malabsorption requiring parenteral nutrition and supplemental fluids to maintain growth and fluid status (4, 6). Patients with MVID are difficult to manage with a number of clinical problems including acidosis, dehydration, central line infections and micronutrient deficiency. Previously often lethal in infancy, advances in parenteral nutrition have now extended life-expectancy for MVID beyond early childhood (7, 8). Despite this, management of MVID is currently primarily supportive and sometimes proceeds to intestinal transplant, with no effective symptomatic or disease modifying therapies.

MYO5B function is critical for normal polarized vesicular trafficking in intestinal epithelial cells and particularly important for protein recycling at the intestinal apical membrane (1, 9–14). Loss of MYO5B function leads to defective microvillus formation and a structurally abnormal apical brush border (1, 11). MYO5B activity is thought to be critical for normal localization of intestinal transport proteins involved in nutrient and fluid absorption and secretion (15–17). Previous studies in MYO5B knockout animal models and intestinal cancer cell lines have shown profound loss or mis-localization of key transporters such as the intestinal sodium-dependent glucose cotransporter (SGLT1) and the sodium/hydrogen exchanger (NHE3) from the apical membrane, with intact cystic fibrosis transmembrane conductance regulator (CFTR) localization (16). Several recent studies examining human MVID intestinal tissues have confirmed these protein changes but the functional balance of fluid absorption *vs* secretion in MVID patient cells remains untested (18). Although animal models and transformed cell studies have been important in our current understanding of MVID pathophysiology, the ability to extrapolate and understand MYO5B-related changes in primary human cells and tissues remains a priority.

Human intestinal stem cell-derived enteroids obtained from endoscopic mucosal biopsy tissue represent a potentially powerful model for understanding protein function, phenotype-genotype correlations, and for therapy development in monogenic epithelial disorders such as MVID (19, 20). Previous studies in genetic disorders affecting the intestine such as Cystic Fibrosis (21) and TTC7A deficiency (22) have highlighted the use of phenotypic and functional assessments for developing and validating therapies and providing patient-specific pre-clinical data. A major barrier for enteroid generation for MVID, however, is that patients rarely undergo endoscopy after initial diagnosis. This along with a need for access to a tertiary medical center with the research resources and logistics required for enteroid generation has led to a lack of availability of human primary enteroids for the study of MVID.

Differentiation and maturation of intestinal epithelial cells from intestinal stem cell-derived progenitor populations into specialized cell types including secretory/sensory cells such as goblet, enteroendocrine and tuft cells as well as mature absorptive enterocytes is dependent on the balance of secreted growth factors that activate Wnt, Notch and TGF-b signaling pathways (23). Previous studies in MVID mouse models have indicated that loss of Myo5b function may lead to altered differentiation and maturation of intestinal cells and that altering intestinal Wnt/Notch balance may be a potential therapeutic strategy (24).

Here, we report the generation and functional investigation of human small intestinal enteroids derived from two patients with MVID due to loss-of-function MYO5B mutations. We show that MVID patient-derived enteroids recapitulate the native epithelial tissue phenotype both structurally and functionally, with loss of microvilli, loss of sodium-dependent electrolyte absorption and intact chloride secretion and fluid secretion. We leverage these functional read-outs of transport function to assess the potential utility of the anti-secretory anti-diarrheal drug Crofelemer for symptomatic therapy for MVID. We further show that loss of MYO5B function leads to defective epithelial differentiation and maturation and that Notch inhibition can structurally and functionally rescue MVID enterocytes. Lastly, we carried out transcriptomic analysis of enteroids following Notch inhibition revealing novel pathways and targets for potential disease-modifying therapies for MVID.

## RESULTS

### Enteroids from MVID patients have a defective brush border and altered differentiation

Duodenal endoscopic biopsies were obtained from two patients with severe intestinal manifestations of MVID (see Supplementary Methods) along with age matched heathy controls and enteroids generated as previously described (25, 26). Patient I was confirmed to have compound heterozygous variants in the MYO5B gene: c.2111delA (p.Phe704Serfs66*) a frameshift mutation and c.1576 G>A (p.Q526*) a paternally inherited novel nonsense mutation that leads to early truncation. These early truncation mutations lead to a functional MYO5B deletion, so the mutations in this patient are referred to as MYO5B KO. Patient II had a confirmed homozygous missense mutation in MYO5B: c.1979C>T (p.P660L), a known missense mutation leading to loss of motor protein function (referred to as P660L in the text)(10).

Confocal and electron microscopy of enteroids (Figure 1A, B and Supplementary Figure 1) indicated loss of apical microvilli and reduced epithelial cell height in MVID cells as compared to a healthy control. MVID enteroids (MYO5B KO and P660L) were grown in standard high Wnt-containing enteroid expansion media (WERN) and passaged similarly to healthy controls, but also showed an increased ability to be expanded and form new enteroids when passaged (Figure 1C, D). When switched to low Wnt differentiation media, healthy control enteroids showed robust differentiation and maturation (Figure 1C, E) with increased expression of markers of mature secretory and absorptive epithelial cells (Muc2, Ngn3, Alpi). In contrast, both MVID enteroid lines had markedly attenuated epithelial marker transcription suggesting that loss of MYO5B function is associated with altered epithelial cell maturation. The appearance of pathognomonic intracellular inclusions was generally rare in the human MVID enteroids and was only noted occasionally in differentiated (ERN) cultures (Supplementary video 1).

**FIGURE 1:**
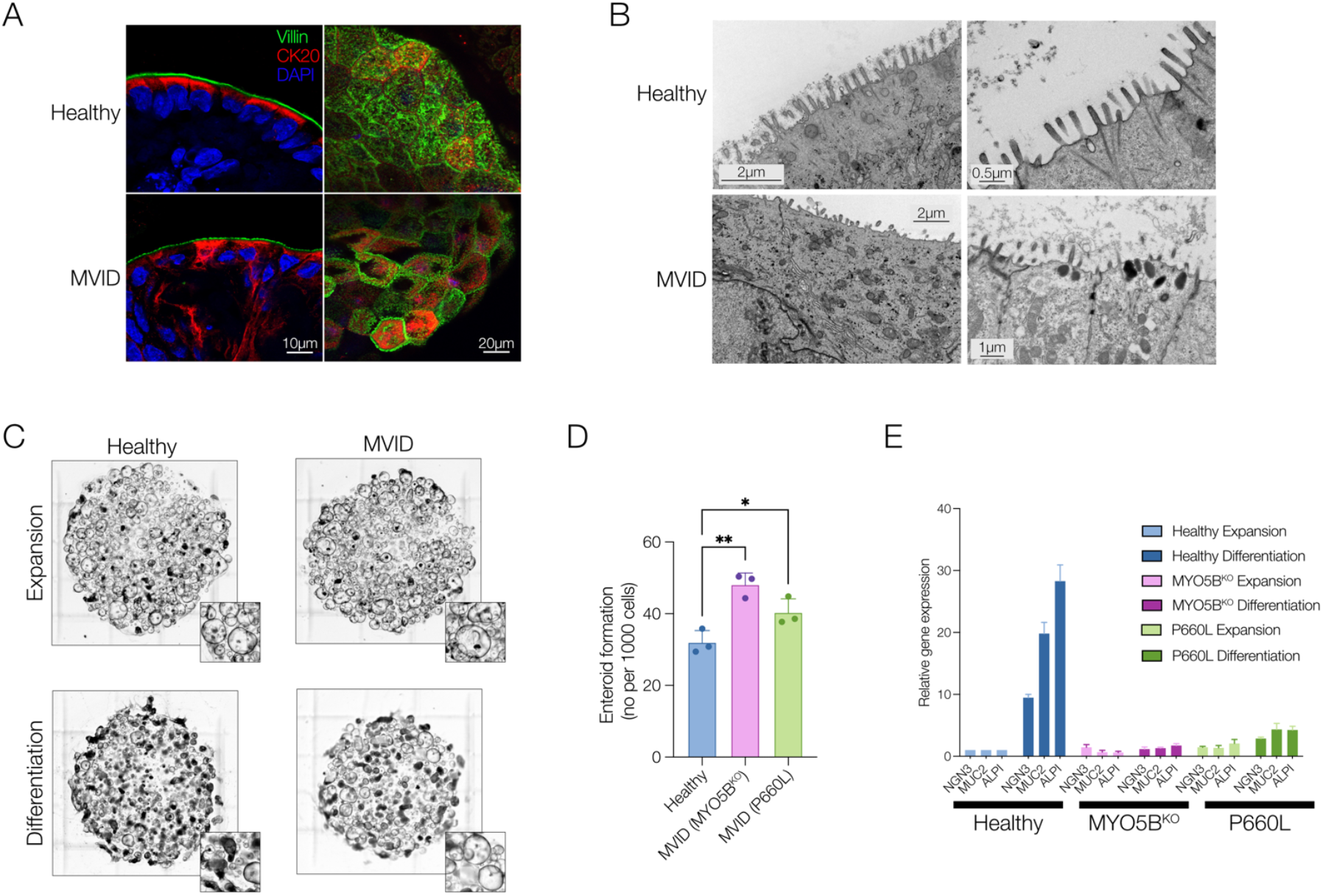
MVID enteroids recapitulate native epithelial disease changes. **A**. Representative confocal images of Villin (green), Cytokeratin 20 (CK20, red) and nuclei (DAPI) of healthy and MVID (MYO5B KO) enteroids in cross-section (left) or en face (right). **B**. Representative electron micrographs of healthy and MVID enteroids. **C**. Brightfield images of enteroid cultures grown in expansion versus differentiation media. Inset images highlighting changes in cultures with increased spheroid (stem-like) morphology in MVID differentiated cultures versus healthy. **D**. Enteroid formation assay showing new enteroid formation (at 4 days post-plating) following enzymatic dissociation and replating. Error bars represent means ± SEM, n=3 experiments *p < 0.05, **p < 0.01. **E**. Relative gene expression (normalized to healthy expansion) for neurogenin 3 (NGN), mucin 2 (MUC2) and alkaline phosphatase (ALPI) in healthy and MVID enteroids following switching to enteroid differentiation media. Error bars represent means ± SEM, n=3 experiments.

### MVID mutations result in altered secretory cell populations

To further assess cell populations and protein localization in MVID, we conducted multiplex immunofluorescence (MxIF) staining (27) and analysis of MVID patient duodenum compared to healthy control tissue. MVID tissues from both patients showed reduced numbers of pEGFR+ tuft cells and DEFA5+ Paneth cells and increased CGA+ enteroendocrine cells (Figure 2A). Apical enterocyte alterations in MVID can be patchy with areas of intact but discontinuous brush border. MVID enteroids (Figure 1A) also exhibited a patchy loss of apical brush border villin staining. To provide quantification of brush border continuity in tissue sections we analyzed staining for CD10, an aminopeptidase and a well-known marker of the brush border in mature enterocytes, by skeletonizing positive signal and measuring line distances (Mean Ferret distance). Using this metric we showed as expected that MVID tissues from both patients show significantly reduced brush border continuity versus controls (Figure 2B).

**FIGURE 2:**
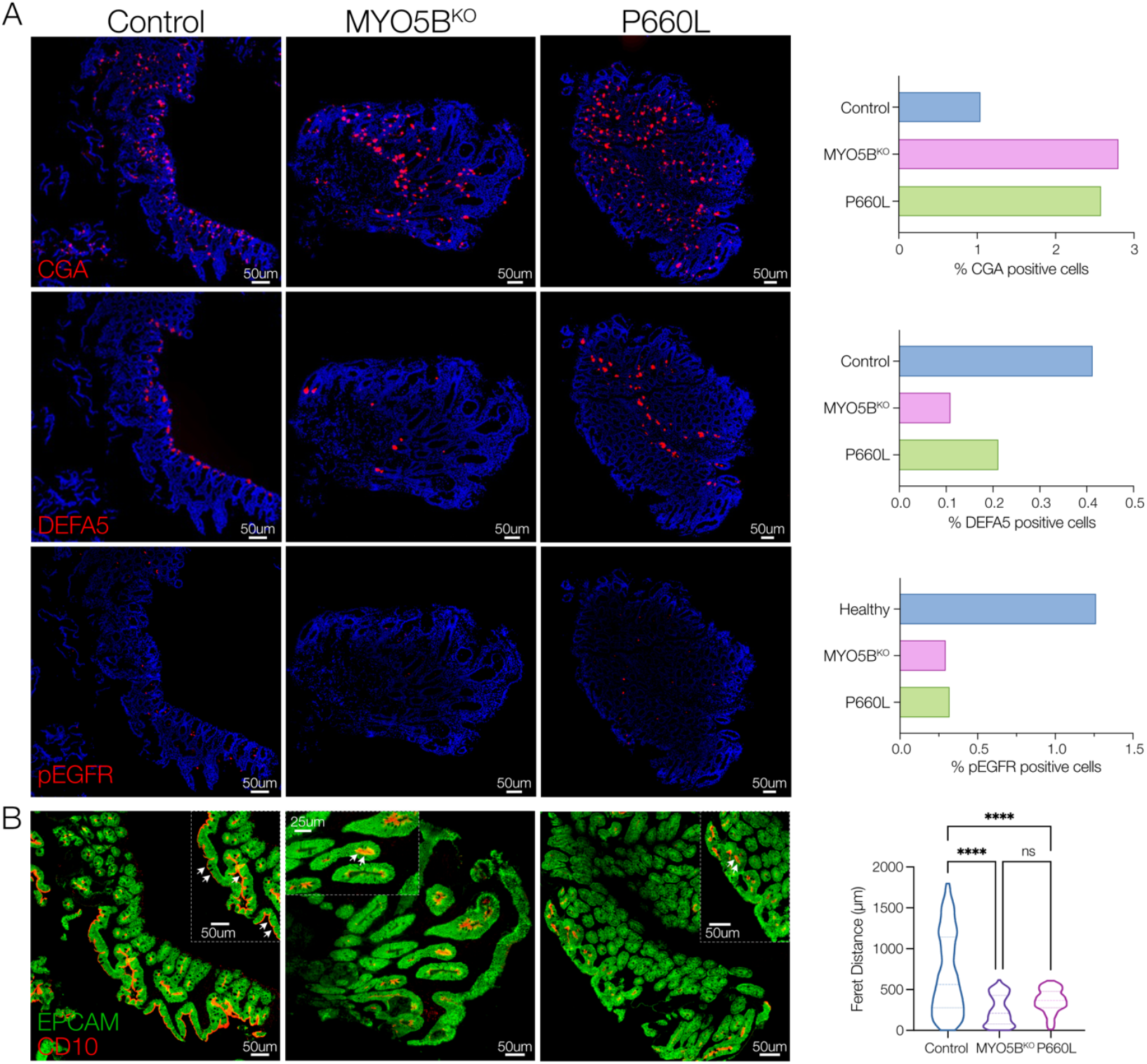
Altered secretory cell populations in MVID patient tissues. **A**. Immunofluorescence images of human duodenal biopsy sections stained for Chromogranin A (CGA), Defensin alpha 5 (DEFA5) and phospho-Epidermal Growth Factor Receptor (pEGFR). Graphs (right) shown whole biopsy counts of positive cells for each antigen. **B**. Representative images of CD10 and Epithelial Cell Adhesion Molecule (EPCAM). Graph (right) of continuity analysis (Feret’s Distance) of linear CD10 staining across all biopsy images.

### Multiplex Immunofluorescence analysis reveals variant-specific alterations in enterocyte transport protein localization in MVID small intestine

A major advantage of MxIF staining is the ability to assess multiple antigens on the same tissue section, allowing cellular protein-protein localization within the resolution of the imaging optics. We applied a targeted panel of antibodies that included epithelial cell type markers, structural markers, and important small intestinal transport proteins (Figure 3A). We conducted a cross-correlation analysis to compare localization of the panel proteins from the two MVID patient tissues against a healthy control (Figure 3B). The overall pattern of correlations between specific pairs of antigens was more like the healthy control for the P660L mutation intestine versus the MYO5B KO. Correlations of membrane-associated transporters with mature enterocyte markers of apical and basolateral membranes were reduced in both MVID patients (NHE3-Villin, and GLUT2-β-catenin). Although SGLT1 showed similar correlations with the apical membrane in healthy and MVID tissues, closer inspection of villus tip enterocytes revealed reduced apical staining and increased internalization in MVID tissues (SGLT1-Villin, Figure 3C). In contrast, NHE3 showed both reduced abundance as well as reduced membrane localization (NHE3-Villin) and significantly altered and mutation-dependent association and localization with MYO5B. These MxIF tissue studies corroborate previous findings of reduced apical transport proteins and alterations in cellular localization in MVID and allow future analysis of genotype-specific changes.

**FIGURE 3:**
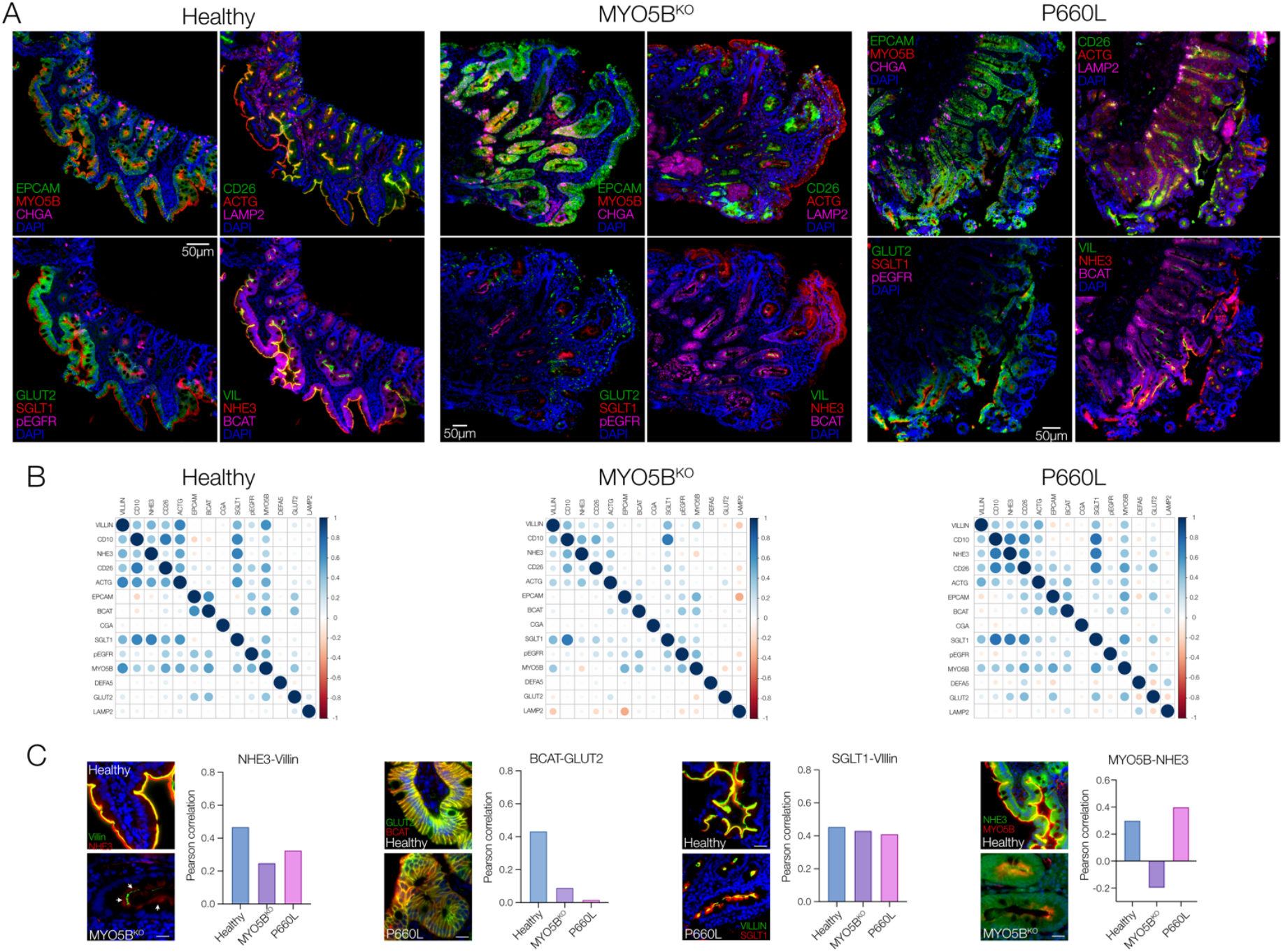
Multiplex immunofluorescence highlights protein localization changes in patient tissues. **A**. Multiplex Immunofluorescence panels representing 12 antigens on duodenal biopsy tissues. **B**. Cross-Correlation Matrices for paired antigens (Pearson’s Coefficient) with dot color indicating direction of correlation and dot size indicating extent of correlation. **C**. Individual pair-wise correlations for Na^+^/H^+^ exchanger 3 (NHE3) and Villin, Beta-catenin (BCAT) and Glucose transporter 2 (GLUT2), Sodium-Glucose Cotransporter 1 (SGLT1) and Villin, Myosin 5b (MYO5B) and NHE3. Inset example images showing staining patterns. Scale bar 10 µm.

### Epithelial sodium-hydrogen exchange is severely impaired, but chloride secretion is intact in human MVID

To functionally assess electrolyte absorption and secretion in human patient MVID epithelial cells, we conducted a series of electrophysiological and fluorometric cellular transport experiments. Patient-derived intestinal cells were cultured as differentiated monolayers on Transwell inserts, and transepithelial short-circuit current (*I*_sc_) was measured using a custom-built multi-well Ussing chamber (Supplementary Methods). MVID cells had a transepithelial electrical resistance similar to healthy controls at the time of measurement (Figure 4A). Phlorizin-sensitive glucose-stimulated *I*_sc_, reflecting SGLT1 transport, was reduced in MVID cells by ∼60% compared to controls (Figure 4B). In contrast, both forskolin (cAMP)- and carbachol (Ca^2+^)-induced chloride secretion was similar between MVID cells and controls (Figure 4C). Chloride secretion stimulated by the muscarinic agonist carbachol (CCh) was notably high in both control and MVID primary duodenal cells. Carbachol-stimulated chloride currents in intestinal cell lines (e.g., T84 cells) are known to be predominantly mediated via the Cystic-Fibrosis Transmembrane Regulator (CFTR). To further assess this, we measured CCh-stimulated currents with and without the CFTR inhibition (CFTRinh-172) (28), in young (2-year-old) healthy and MVID duodenal cells versus cells derived from an adult grown in the same conditions. Interestingly, the contribution of non-CFTR, most likely calcium-activated chloride channel (CaCC), was lower in adult duodenal cells than either early childhood duodenal cells, suggesting important age-dependent differences.

**FIGURE 4:**
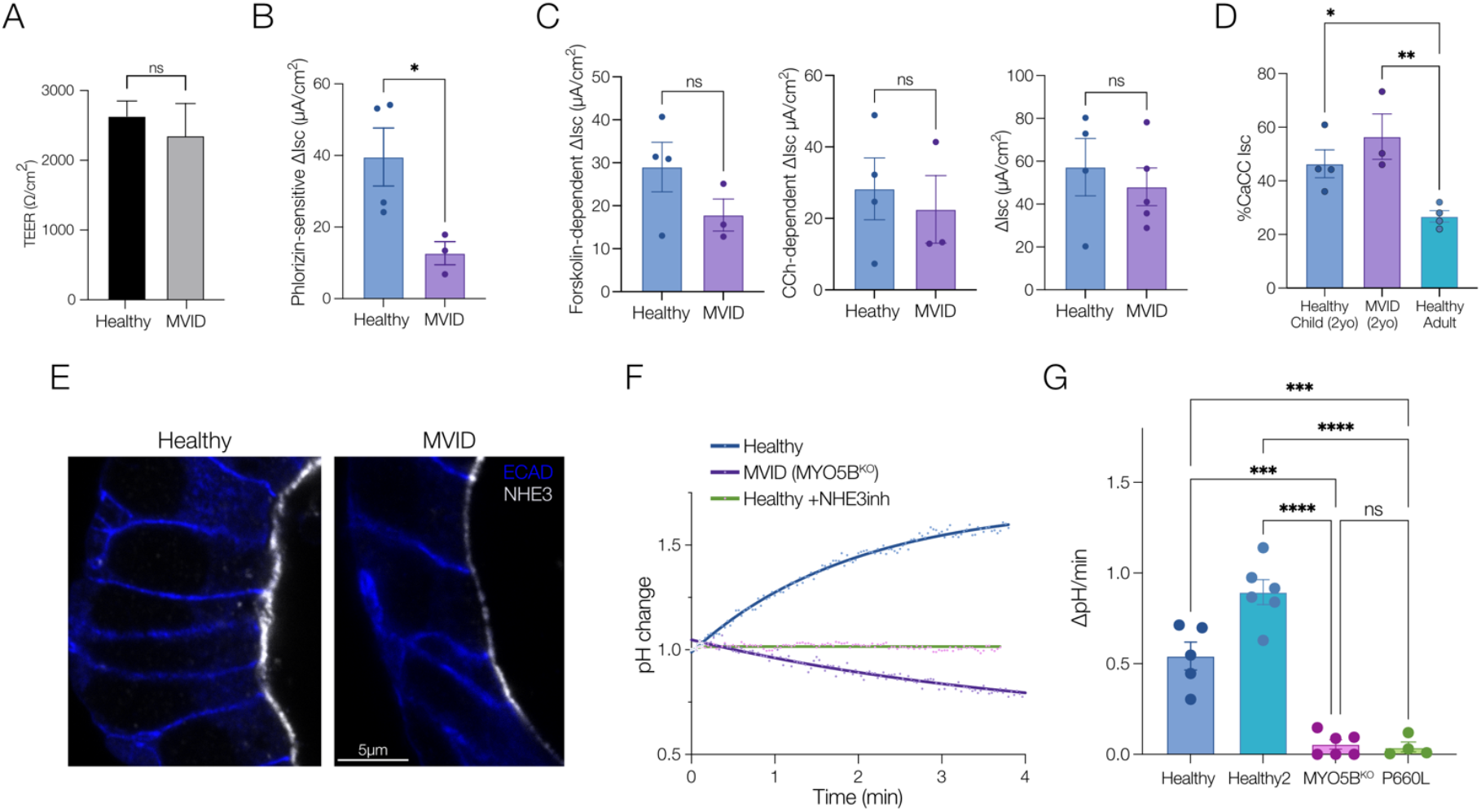
Loss of sodium absorption and normal chloride secretion in MVID patient enteroids. **A**. Transepithelial Resistance (TEER) on 10-14 days post plating on Transwell inserts for short-circuit current measurements. **B**. Glucose (20 mM)-stimulated, phlorizin-inhibitable short-circuit current (Isc) in healthy and MVID (MYO5B KO) monolayers. Error bars represent means ± SEM, n=3-4 experiments, *p < 0.05. **C**. Forkolin (10 µM) stimulated Isc. Error bars represent means ± SEM, n=3-4 experiments (left), Carbachol (CCh, 50 µM) stimulated Isc. Error bars represent means ± SEM, n=3-4 experiments (middle), Combined forskolin and Carbachol stimulated Isc. Error bars represent means ± SEM, n=3-4 experiments (right). **D**. % Carbachol (CCh, 50 µM) stimulated Isc not inhibited by CFTR-inh-172 (50 µM) in healthy control cells grown from a young donor (2yo), adult donor (20yo) and MVID patient (MYO5B KO). **E**. Super-resolution images (STED) of NHE3 localization and abundance in healthy and MVID patient enteroids. **F**. Example curves showing change in pH calculated from analysis of intensity changes of SNARF-5F fluorescence in healthy and MVID cells and healthy cells after pre-treatment with the NHE3 inhibitor (10µM). **G**. Summary graph showing NHE3-dependent pH changes in healthy and MVID patient cells. Error bars represent means ± SEM, n=4-6 experiments, ***p < 0.001, ***p < 0.0001.

Similar to MxIF tissue staining, confocal imaging of enteroids showed markedly reduced localization of NHE3 at the apical membrane of MVID cells. To functionally assess electroneutral Na^+^/H^+^ exchange, enteroids were grown on glass coverslips for fluorescence imaging. Cells were loaded with the intracellular pH-sensitive fluorophore SNARF-5F and placed in an imaging chamber. Using a previously optimized Na^+^ free alkalinization protocol, the kinetics of intracellular pH changes as a measure of NHE3 activity was monitored in patient-derived duodenal cells (29). As shown in Figure 4F, healthy control cells exhibited a robust change in pH that was blocked by pre-treatment with an NHE3 inhibitor (S3226). MVID cells from both patients showed reduced and, in some experiments, complete loss of NHE3 activity (Figure 4G).

### An approved natural-compound anti-diarrheal can inhibit chloride and fluid secretion in MVID epithelium

MVID patients can have very large daily fluid loss (2–5 L per day), with consequent significant requirement for fluid replacement and a tenuous acid/base status. Our functional assessment of MVID cell transport showed severe loss of Na+ mediated fluid absorption in the setting of potentially intact chloride-mediated fluid secretion, suggesting that blocking chloride transport may theoretically improve fluid balance in MVID. To further assess this, we obtained the FDA approved anti-diarrheal compound, Crofelemer, to test in MVID patient cells. Crofelemer is a proanthocyanidin, natural anti-diarrheal reagent that inhibits both the CFTR- and CaCC- mediated chloride secretion (30). Analysis of combined forskolin- and carbachol-stimulated *I*_sc_ showed a robust and dose-dependent inhibition with an IC_50_ ∼ 30 µM (Figure 5A and 5B), similar to previous reports, but with an increased maximal inhibition compared to previous data in colonic cell lines (∼80% vs 50%) (Figure 5C) (30). To measure fluid secretion, we employed an 3-dimensional enteroid swelling assay (21, 22) following stimulation of enteroid fluid secretion with forskolin and tested the efficacy of Crofelemer. Corroborating our chloride current data, both healthy and MVID enteroids exhibited a robust induction of fluid secretion, which was inhibited by approximately 40% and 50%, respectively, by administration of Crofelemer (100 µM) and inhibition continued up to 4 hours post-stimulation (Figure 5F).

**FIGURE 5:**
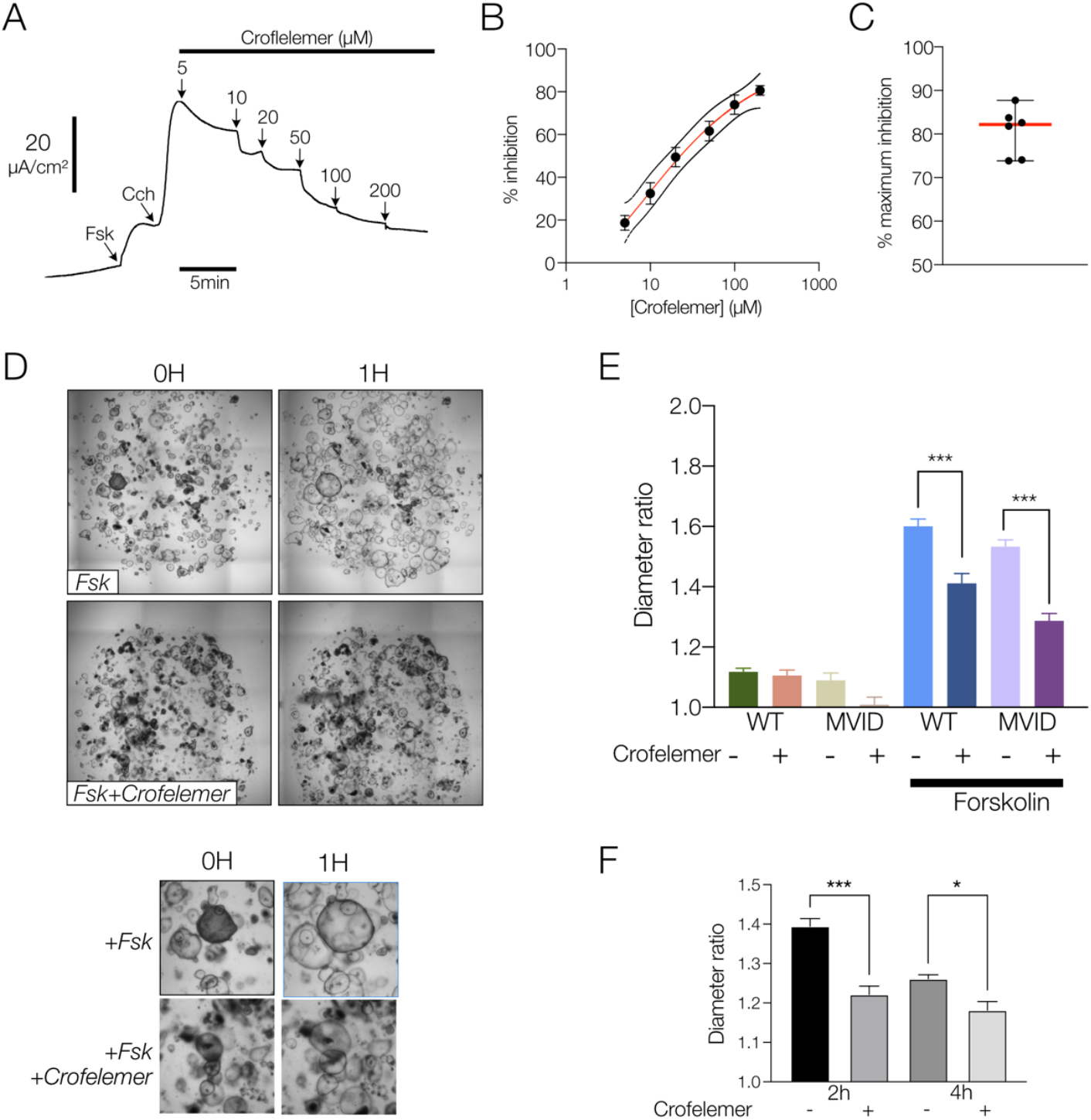
Crofelemer inhibits chloride and fluid secretion in MVID patient enteroids. **A**. Representative curve showing dose dependent inhibition of Crofelemer on forskolin (10 µM)- and carbachol (CCh, 50µM)-stimulated Isc. **B**. Dose-response curve for Crofelemer-induced inhibition of forskolin and carbachol stimulated Isc in MVID patient enteroids (MYO5B KO). Error bars represent means ± SEM, n=6 experiments **C**. Maximal percent inhibition of agonist stimulated current by Crofelemer. Error bars represent means ± SEM, n=6 experiments. **D**. Example brightfield images before and after forskolin (2 µM) ± Crofelemer (200 µM) in MVID patient enteroids. Inset below showing enteroid swelling. **E**. Summary graph showing increase in enteroid size (diameter ratio) in healthy and MVID enteroids at 1 hour ± forskolin and ± Crofelemer (200µM). Error bars represent means ± SEM, >300 enteroids from 4 experiments. **F**. Diameter ratio in MVID enteroids at 2 and 4 hours post stimulation ± Crofelemer (200µM). Error bars represent means ± SEM n=3 experiments.

### Notch inhibition can structurally and functionally correct MVID epithelial defects

Although attenuation of fluid losses would be of benefit for MVID patients, restoration of normal epithelial function including fluid absorption following MYO5B loss of function would have major implications for disease management. Previous studies in mouse models (16, 24) as well as our finding of impaired differentiation in patient enteroids, indicated that alteration of Wnt/Notch signaling may allow functional epithelial recovery. Treatment of healthy enteroids with the γ-secretase inhibitor, DAPT, induced robust differentiation and polarization of cells (Figure 6A, Supplementary Figure 1, 2). DAPT-treated MVID enteroids remarkably showed recovery of microvillus structure, with increased microvillus length (Figure 6B) and length of the actin core (Figure 6C). Overall cell polarization of MVID cells as measured by the organelle free zone at the apical aspect of columnar epithelial cells was also increased following Notch inhibition (Figure 6D). In addition to structural recovery, NHE3 abundance at the brush border was increased (Figure 6E) and there was significant recovery of NHE3 function as measured by direct Na^+^/H^+^ exchange activity of MVID cells from both patients (Figure 6F, 6G).

**FIGURE 6:**
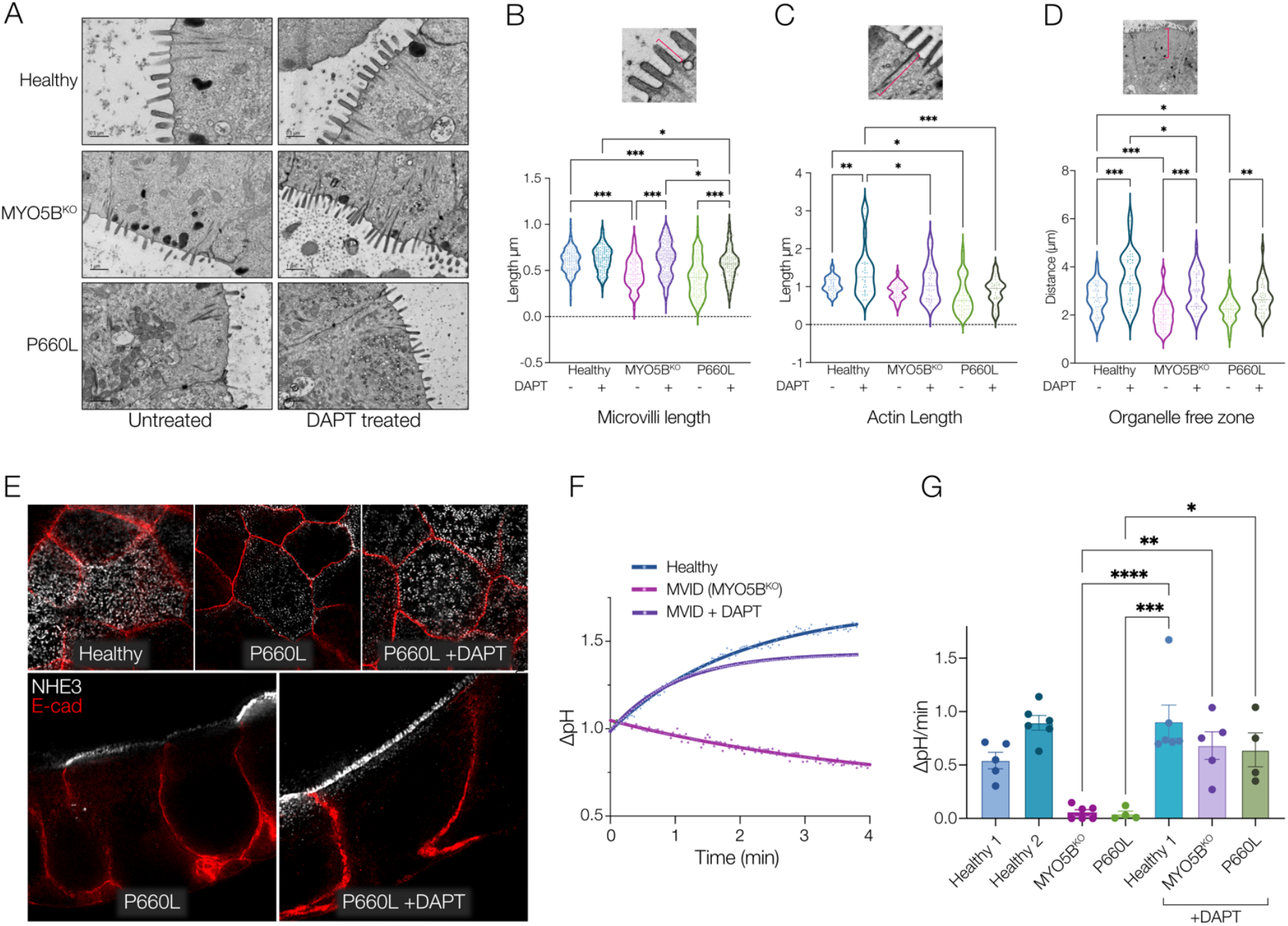
Notch inhibition rescues MVID patient enteroid differentiation. **A**. Electron micrographs of healthy and MVID enteroids ± DAPT treatment (10 µM). **B**. Analysis of EM images of microvillus length. **C**. sub-apical actin bundle length. **D**. Distance of apical organelle free zone. Inset images above show example measured parameter. Graphs show measurements from at least 10 EM images. **E**. Super-resolution confocal images (STED) en-face (top) and cross-section (below) of NHE3 localization and abundance in healthy and MVID patient enteroids following treatment with DAPT (10 µM). Representative curves showing change in intracellular pH in healthy cells and MVID cells ± DAPT **G**. Summary graph showing NHE3-dependent pH changes in healthy and MVID patient cells ± DAPT. Error bars represent means ± SEM, n=4-6 experiments, *p < 0.05, **p < 0.01, ***p < 0.001, ***p < 0.0001.

### Transcriptomic analysis of Notch inhibition in MVID enteroids reveals a novel target pathway for MVID therapy

Given the evidence for functional and structural recovery in MVID cells with different mutations following Notch inhibition, we sought to identify potential pathways and target proteins that may mediate the effect. We therefore conducted unbiased bulk RNA sequencing of healthy versus MVID enteroids (MYO5B KO) with and without DAPT treatment. Figure 7A-C shows the large number of up- and down-regulated genes associated with DAPT treatment as well as MYO5B loss of function (Supplementary file 1). To detect genes of interest, we identified genes (Figure 7D, dotted boxes) that were significantly altered at baseline between MVID and healthy enteroids and that were significantly altered in the opposite direction following DAPT treatment (e.g. downregulated in MVID at baseline and upregulated following DAPT treatment). To further prioritize, we filtered these hits based on log2Fold Change (FC), False Discovery Rate and base mean (Figure 7E). Gene ontology and protein interaction analysis (Figure 7F, Supplementary Figure 3) implicated genes involved in transport regulatory activity, metabolic processes, and interestingly, PPARα signaling. The top upregulated gene hits in our prioritized analysis included *RAB32, SGK2, PDZK1, ANPEP* and *ANXA13*. ANPEP is an apical membrane aminopeptidase and ANXA13 is an annexin family member associated with the differentiated villus enterocytes, and therefore we reasoned that these likely reflected recovery of the brush border in keeping with our structural analysis. SGK2, RAB32 and PDZK1 have been implicated in regulation of trafficking and/or alteration of membrane transport function. We therefore validated these changes by probe specific qPCR analysis (Figure 8A). The activity of Serum/Glucocorticoid Regulated Kinase (SGK) family of enzymes has been previously implicated in MVID and MYO5B function (17). Immunoblot of SGK2 protein (Figure 8B) indicated a DAPT-induced increase in SGK2 in both MVID patients’ cells consistent with the RNA expression data. To functionally assess whether SGK2 may be a potential target for recovery of MYO5B function, we tested NHE3 activity. DAPT-mediated increases in NHE3 activity in MVID cells were inhibited by pre-treatment with an SGK inhibitor, suggesting that SGK function may play a mechanistic role in DAPT-mediated epithelial recovery in MVID.

**FIGURE 7:**
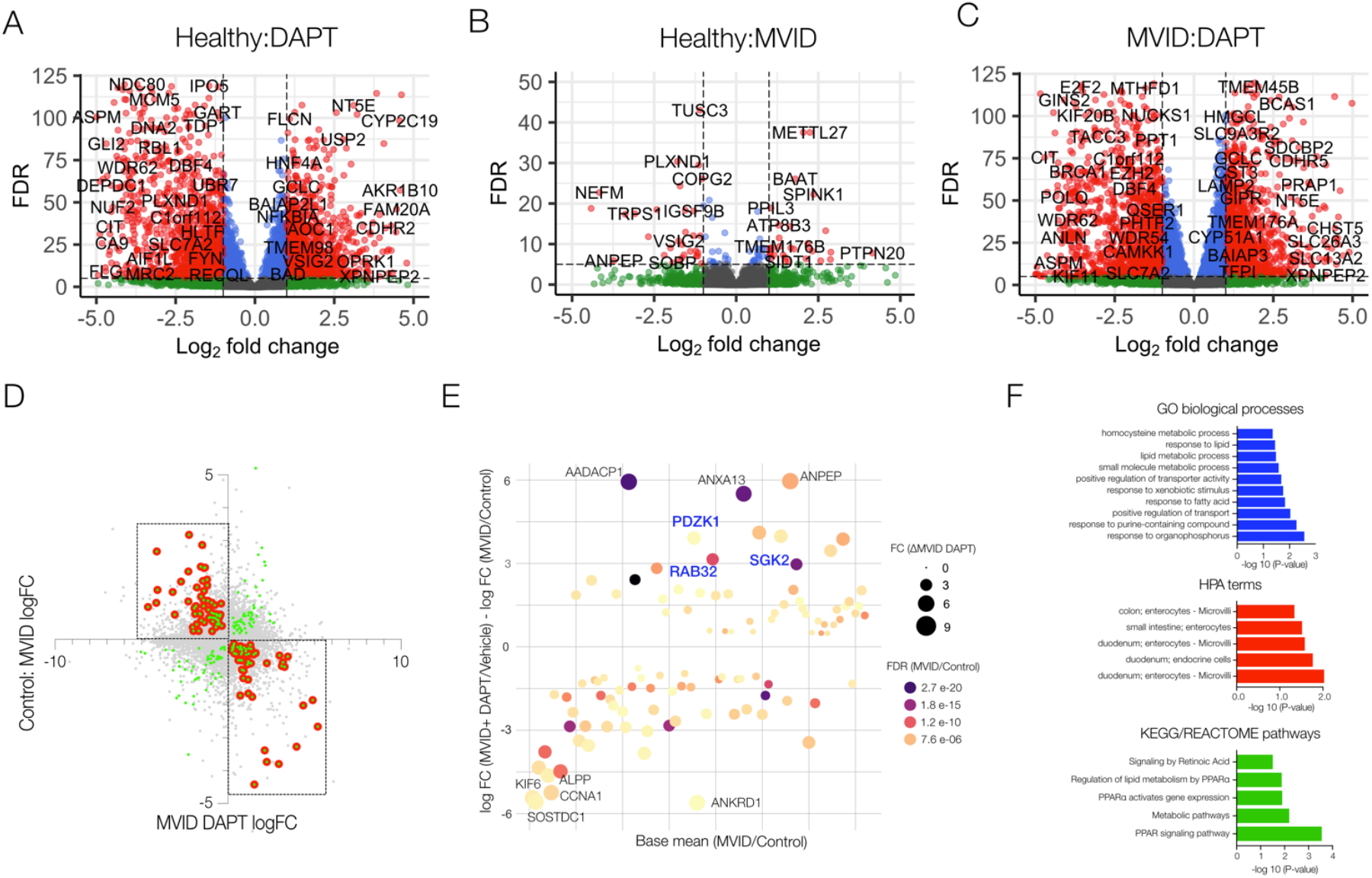
Genome-wide transcriptomic analysis reveals potential targets for rescue of MVID enteroids. **A**. Volcano plot showing log_2_ fold change and false discovery rate (FDR) showing genes with significantly up- and down-regulated expression (red) in healthy enteroids (n=3) following DAPT treatment (10 µM). **B**. Volcano plot showing genes with significantly up- and down-regulated expression (red) between healthy enteroids and MVID enteroids (MYO5B KO) (n=3). **C**. Volcano plot showing genes with significantly up- and down-regulated expression (red) in MVID enteroids (MYO5B KO) following DAPT treatment (10 µM). **D**. Plot of genes with significantly altered expression (green dots) between MVID and healthy against MVID + DAPT. Red dots indicate genes changing in opposite directions following DAPT treatment (filtered genes). **E**. Dot plot of filtered genes by change in expression and base mean expression, with dot size indicating fold change and color indicating FDR. Highlighted genes based on previous functional data indicating plausible biological role. **F**. Pathway analysis showing most significant GO terms, HPA terms and KEGG pathways.

**FIGURE 8:**
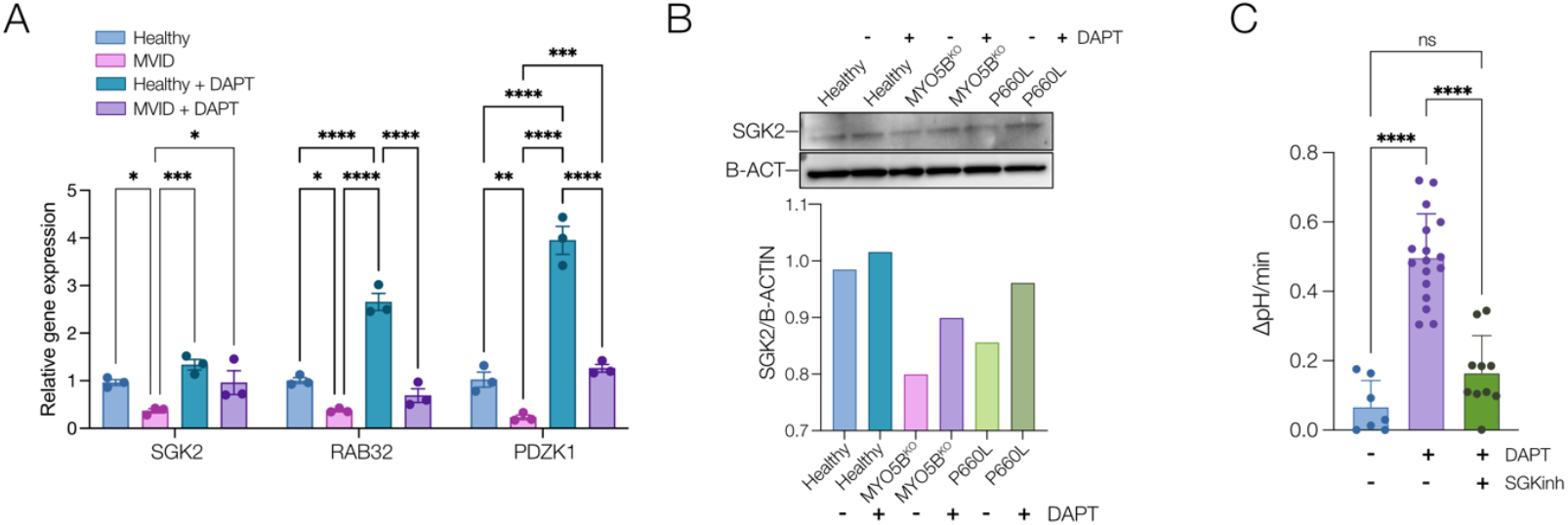
Analysis of potential genes involved in DAPT-mediated rescue of MVID enteroids. **A**. qPCR analysis showing relative gene expression (normalized to healthy untreated) for Serum- and glucocorticoid-induced protein kinase 2 (SGK2), Rab GTPase 32 (RAB32) and PDZ domain-containing 1 (PDZK1) in healthy and MVID enteroids ± DAPT (10 µM). Error bars represent means ± SEM, n=3 experiments. **B**. Immunoblot (top) of SGK2 protein changes ± DAPT with densitometric quantification (below) normalized to beta-actin. **C**. Summary graph showing NHE3-dependent pH changes in healthy and MVID patient enteroids ± DAPT (10 µM) or MVID patient enteroids ± DAPT and SGK inhibitor (GSK650394, 5 µM). Error bars represent means ± SEM, ****p < 0.0001.

### DISCUSSION

Intestinal epithelial MYO5B is a critical regulator of epithelial polarity and the establishment of apical membrane recycling and apical trafficking (1, 3, 11, 12). Loss of MYO5B function, associated with MVID, results in a structurally abnormal brush border in patients and mouse models, with decreased localization of brush border transporters important for intestinal fluid and nutrient absorption and homeostasis. Enteroids and tissue derived from both MVID patients described here showed broadly similar phenotypic features found in previous descriptions of MVID patient tissues, with loss of microvilli and loss of transport protein localization (16, 18). Our MxIF method allowed further analysis of protein localization and identification of differences in localization patterns in tissue between the two patients reflective of the impact of different MYO5B mutations. Previous studies have shown that the P660L missense mutation induces a rigor status in MYO5B (10). The second MVID tissue and enteroids reported here possess two severe truncating mutations, which likely lead to deletion of MYO5B.

In addition to structural and localization changes, we directly investigated transport function in MVID and healthy epithelial cells. There is minimal data on electrolyte transport function in patient-derived human enteroids and existing data has been restricted to adult samples (29). We found reduced SGLT1 dependent Na^+^ transport and near complete loss of NHE3 transport with normal chloride secretion in MVID cells. Previous studies in MYO5B KO Caco-2 cells, a pig model of MYO5B(P660L) and MYO5B KO mice have suggested maintenance of chloride and therefore fluid secretion in MVID (15, 17, 18). Our data corroborate these findings and suggest that loss of Na^+^ mediated transport, particularly via NHE3, may underlie the majority of the fluid loss. Quantitatively most of the fluid (5L/day in an adult) required to be absorbed (primarily by Na^+^ dependent mechanisms) in the small intestine is generated endogenously by the gastrointestinal tract. Strategies to enhance or prevent loss of NHE3 function may therefore have considerable benefit for therapy in MVID. An interesting finding was the increased proportion of putatively calcium-activated chloride channel vs CFTR activity in young children likely reflecting developmental differences in duodenal transport. Further studies to carefully characterize factors such as developmental age, sex and environmental variables in healthy cells on transport and other epithelial functions are needed to provide normative data to understand changes seen in disease states affecting the proximal small intestine including celiac disease, inflammatory bowel disease and environmental enteric dysfunction.

The maintenance of normal chloride and fluid secretion in the setting of loss of Na+ transport likely further exacerbates fluid loss at baseline, but particularly during periods of stress such as infection where chloride channel secretion is stimulated. To assess a potential immediate therapeutic intervention, we conducted pre-clinical studies using the approved anti-diarrheal Crofelemer, a combined CFTR and calcium-activated chloride channel blocker. Short-circuit current analysis showed robust inhibition of stimulated chloride currents with a higher maximal inhibition than previously reported. Previous data in T84 cells suggest that Crofelemer has increased potency on the putative calcium-activated chloride channel vs CFTR (30) and the age-dependent differences we found in these currents may underlie this finding. If so, Crofelemer would be predicted to be more effective in young children than adults. Crofelemer was also able to inhibit maximally stimulated fluid secretion significantly, but incompletely in enteroids. Fluid management of MVID patients is a prominent clinical issue and even a modest (10-20%) decrease in intestinal fluid output would translate to a significant symptomatic improvement. These data therefore suggest that inhibition of chloride secretion may be a viable strategy for symptomatic management in MVID.

We found that enteroids exhibited altered differentiation characteristics versus healthy controls consistent with previous mouse MYO5B KO data showing disrupted stem cell marker expression, a loss of tuft cells, and MYO5B loss-associated decreased transcription levels of Wnt ligands with maintained expression of Notch signaling molecules (24). The balance of proliferation and differentiation in the intestinal epithelium is a complex process that is driven by the balance of stem cell activity and relative specification to the various mature cell types (secretory cell vs enterocyte) and governed in tissue by the balance of niche growth factor signals (23). Enteroid models are subsequently an inherently regenerative model where the exogenous supplementation of growth factors can only approximate the endogenous tissue state. Therefore, functional and transcriptomic data should be interpreted in the setting of the specific growth conditions present in a particular experiment. However, an advantage of the system is the ability to directly manipulate growth factors to direct epithelial cell state and compare changes. Given the previous data suggesting imbalanced Wnt/Notch signaling with MYO5B loss and the enhanced epithelial differentiation by a gamma-secretase inhibitor, DBZ, in MYO5B KO mouse model (24), we used Notch inhibition to test the possibility of functional recovery in patient enterocytes. Notch inhibition via DAPT induced significant increases toward differentiation in all cell lineages in healthy enteroids and partially recovered the defective differentiation seen in MVID enteroids grown in standard differentiation media. Notably, DAPT-treated MVID epithelial cells showed structural evidence of increased polarization and maturation with increases in cell height, microvillus height, and brush border expression of NHE3. We have demonstrated in MVID patient-derived enteroids that the epithelial defects in MVID may be reversible and that loss of MYO5B function can be bypassed by targeting the pathways involved in Wnt/Notch signaling.

To understand the basis of DAPT mediated rescue in MVID potential pathways, we carried out transcriptomic analysis of DAPT-treated enteroids. DAPT treatment alone on healthy enteroids caused a shift in the transcriptomic landscape with over 8000 genes significantly changed. We narrowed down potential candidate genes by focusing on genes altered in MVID versus healthy enteroids and subsequently changed in the opposite direction by addition of DAPT. Using this strategy, we identified several potential genes that may be involved in the functional recovery and further interrogation of this dataset will likely provide insights for both Notch-dependent and MYO5B-dependent intestinal epithelial processes. We focused on several genes that may be plausibly involved in cellular pathways related to MYO5B, based on previous data and known functional roles.

An interesting target identified was Serum- and glucocorticoid-induced protein kinase 2 (SGK2) which was downregulated in MVID patient enteroids relative to healthy controls, but significantly upregulated upon DAPT treatment. SGK2 is part of a family of closely homologous kinases (SGK1-3) involved in cell stress responses (31, 32). SGK function in epithelial cells is thought to be involved in regulation of apical electrolyte transport, with evidence of regulation of sodium re-absorption in the kidney and direct regulation of NHE3 activity via NHE3 regulatory factor (NHERF) proteins (33, 34). SGK expression is regulated by many transcription factors including AP-1, NF-κB, GATA, Ets-2, and p53, and there is one report relating to Notch signaling in macrophages (35–38). In kidney, SGK2 but not SGK1 regulates NHE3 expression and activity by altering the stability of the transporter at the apical membrane (34, 39). In terms of linking SGK function and MYO5B, there is evidence that regulation of SGK1 activity can be mediated by interaction with the NHERF complex via phosphoinositide-dependent protein kinase 1 (PDPK1) linking SGK function with activity and localization of apical transport (40). PDPK1 has been directly linked with MYO5B function with studies suggesting that PDPK1-dependent signaling may provide a therapeutic target for treating MVID (17). Given the suggestive evidence in the literature, our findings that both expression and protein abundance of SGK2 is upregulated by DAPT, and that inhibition of SGK activity can block rescue of NHE3 function provides some preliminary mechanistic data of SGK2 involvement in bypassing the effects of MYO5B loss-of-function.

Other candidates potentially involved in regulation of apical trafficking and protein localization and identified in our transcriptomic screen include Rab GTPase 32 (RAB32) and PDZ Domain Containing 1 (PDZK1). Rab32 is multi-faceted regulator linked to multiple cellular functions including endosomal trafficking, mitochondrial dynamics, and biogenesis of lysosome-related organelles (41–43). Rab32 regulates these cellular pathways through interacting proteins that include the AP-1 and AP-3 adaptor protein complexes and may provide an alternative vesicular pathway for apical trafficking. PDZK1 also known as NHE3 regulatory factor 3 (NHERF3) is a PDZ-domain scaffolding protein that mediates plasma membrane localization and regulation of proteins, and most notably is part of the regulatory complex for NHE3. PDZK1 is thought to form heterodimers with other NHERF family members (NHERF1 and 2) and plays a role in regulation of NHE3 localization and function as well as formation and maintenance of the microvillus structure (44–46). Our studies therefore reveal several lead targets including SGK2 and PDZK1 that can form the basis for future studies of the specific mechanism(s) involved in Wnt/Notch rebalancing-based rescue in MVID and for future targeting of therapies.

In summary, our studies show that patient-derived enteroids recapitulate the cellular phenotype of MVID epithelial cells and provide functional and structural data in human patient cells including identifying key ion and fluid transport changes present in MVID. We leveraged this unique resource to conduct therapeutic testing for symptomatic management and show that therapeutic modalities can bypass trafficking blockades induced by MYO5B loss of function.

## MATERIALS AND METHODS

### Human Subjects Approval

The institutional review board of Boston Children’s Hospital and Phoenix Children’s Hospital approved this study (BCH IRB P00027983, PC IRB 10-019) and informed consent/assent was given in accordance with the Declaration of Helsinki.

### Whole-exome and Sanger sequencing (WES)

DNA from patient 1 and mother were sent to GeneDx for whole-exome sequencing (WES)^9^.(PMID: 32655885). A sequencing library was prepared using the Illumina Exome Enrichment Protocol and captured libraries were sequenced on Illumina HiSeq 2000 or 4000. Sequences were aligned to the human genome reference sequence (hg19). The sequences were aligned to the reference sequence using BWA (Burrows-Wheeler Aligner, version 0.7.15) and variants called with Gatk best practices (version 3.7). The data were filtered to include variants with an allele frequency of <0.001 in publicly available normative databases (gnomAD). Variants were identified using the NextCode Genuity platform. DNA from patient 2 was sent for whole-exome sequencing via a targeted gene panel. Physiochemical differences between the canonical and patient amino acid sequences were determined using CADD, and PolyPhen-2, MutationTaster and SIFT were also used to estimate the impact of the variant on DNA and protein level. The identified *MYO5B* variants was confirmed by Sanger sequencing. The relevant portion of the gene was PCR amplified and Sanger sequencing was performed. The bi-directional sequence was assembled, aligned to reference gene sequences based on human genome build GRCh37/UCSC hg19 and analyzed for known familial sequence variant(s). Sequence alterations were reported according to the Human Genome Variation Society (HGVS) nomenclature guidelines.

### Materials and Reagents

Forskolin and carbachol were purchased from Sigma Aldrich. Crofelemer drug substance was provided by Napo Pharmaceuticals. Antibody for SGK2 was purchased from Cell Signaling (SGK2 (D7G1) Rabbit mAb Catalog # 7499S). GSK650394, the inhibitor against SGK2 was purchased from Selleck Chemicals (Catalog No.S7209). SNARF-5F (and -6) Carbolic acid AM ester, acetate was purchased from ThermoFisher (Cat: S23923) and CFTRinh-172 was purchased from Tocris (Minneapolis, MN).

### Enteroid Culture

Duodenal biopsies were obtained and cultured using methods modified from Sato et al (25). Briefly, crypts were dissociated from duodenal biopsies obtained from patients with MYO5B mutation or from an age-matched healthy control patient. Isolated crypts were suspended in Growth Factor Reduced Phenol Red Free Matrigel (Corning, NY) and plated as 50 μl domes in a tissue culture-treated 24-well plate (Thermofisher) with growth factor (Wnt, R-spondin, Noggin) supplemented media (See Supplemental Methods for media composition and detailed culture methods). Enteroid cultures were passaged by removal of Matrigel with Cell Recovery Solution (Corning, NY), mechanical dissociation of enteroids, and replating in Matrigel every 4 days.

For electrophysiological assessments, enteroids were grown on collagen-coated 0.33-cm^2^ transwell inserts (Costar Corning, CLS3472) and incubated in 95% O_2_/5% CO_2_ at 37°C for at least 7 days. Medium was changed every 3-4 days. Transepithelial electrical resistance (TEER) was measured using an epithelial volt/ohm meter (EVOM; World Precision Instruments).

### Electron Microscopy

Enteroids were embedded in low melting point sea plaque agarose (Cambrex Bio Science, Rockland, ME) creating faux tissue blocks. Samples were then immersion fixed in 2.5% glutaraldehyde (Electron Microscopy Sciences, Hatfield, PA), 2% formaldehyde (Electron Microscopy Sciences) in 0.1M Sodium Cacodylate (Sigma-Aldrich, Burlington, MA) at pH 7.4 for at least 1 hr at room temperature and then at 4°C overnight. Samples were washed with 0.1 M Sodium Cacodylate, and then post-fixed for 1 hr at 4°C in 1% osmium tetroxide (Electron Microscopy Sciences) in 0.1 M Sodium Cacodylate. Samples were then washed in DI water and incubated in 2% aqueous uranyl acetate (Electron Microscopy Sciences) overnight at 4°C. The following day, samples were washed with DI water and then dehydrated at 4°C in a graded ethanol series. The agar blocks were then brought to room temperature and dehydrated with 100% ethanol (Sigma-Aldrich) followed by propylene oxide (Electron Microscopy Sciences). Infiltration in LX112 resin (Ladd Research Industries, Williston, VT), was followed by embedding in flat bottom Beem capsules (Electron Microscopy Sciences). The resulting blocks were sectioned using a Leica Ultracut E ultramicrotome (Leica Microsystems, Wetzlar, Germany) and the sections were placed on formvar (Electron Microscopy Sciences) and carbon-coated grids. The sections were contrast stained with 2% uranyl acetate followed by lead citrate (Sigma-Aldrich), and imaged in a JEOL 1400 transmission electron microscope (JEOL, Peabody, MA) equipped with a Gatan Orius SC1000 digital CCD camera (Gatan, Pleasanton, CA).

### Confocal, STED and Lightsheet Imaging

Confocal imaging was conducted using a Zeiss 880 Airyscan microscope with Zen Black and Blue for control and analysis (Zeiss Instruments, Jena, Germany). STED imaging was conducted using a Abberior StedyCon system (Abberior Instruments GmbH, Göttingen, Germany) with post-image deconvolution using Huygens Pro (Scientific Volume Imaging, Hilversum, Netherlands). Lightsheet imaging was conducted using an ASI DiSPIM microscope (Applied Scientific Instruments, Eugene, OR).

### Multiplex Immunofluorescence

FFPE slides were deparaffinized and incubated in Trilogy® (Sigma) for antigen retrieval. The slides were cover-slipped with 50% glycerol in 0.1 M PBS containing 1 μM Hoechst 33342, and whole slide images were scanned using a Leica/Aperio Versa 200 with 20x objective (Leica Biosystems, Buffalo Grove, IL) for acquisition of autofluorescence signals of the tissues. Coverslips were gently removed, and the slides were pre-blocked with Dako serum-free protein blocking solution (X0909) for 1 hour at room temperature (r/t) or overnight at 4°C. Primary conjugated antibodies listed in Supplementary Methods were applied to the slides for 1 hour at room temperature. Some antibodies were labeled with Zenon rabbit IgG labeling kits (Invitrogen) according to the manufacture’s instruction. Fluorescence signals were quenched by carbonate buffer (pH 11) immediately after scanning of staining signals. Background signals were imaged after each quenching step and subtracted from following staining images. The MxIF data generated in this study are available at PediCODE (COngenital Diarrhea and Enteropathy) Consortium website.

### Multiplex Imaging analysis

Background corrected immunofluorescence image were used for all multiplex image analysis.

#### Percentage of enteroendocrine and tuft cells

These set of analysis was carried out using FIJI ImageJ. The DAPI channel was first histogram normalized and the number of nuclei was determined by the “Find Maxima” function with a prominence parameter of 15. Analogously, the number of endocrine and tuft cells were determined using the channels for pEGFR and CGA respectively with a prominence factor of 40 and 100 respectively.

#### Measuring Feret’s distance

We used a home-written Matlab code for this analysis. The CD10 was first thresholded, binarized and finally skeletonized with a minimal branch length of 10 pixel to avoid background. The maximum caliper of the Feret distance for each of the skeleton in the image was calculated.

#### Measuring correlation between all channels

The correlation coefficient for each channel with every other channel was calculated using Matlab. The correlation matrix was imported into R and hierarchically clustered and plotted using the Corrplot library.

### Short-circuit current measurement

Following the formation of a monolayered enteroids, medium was removed, and the cells were rinsed and bathed in buffer solution (in mM) (130 NaCl, 0.47 KCl, 0.124 MgSO_4_, 0.33 CaCl_2_, 10 HEPES, 2.5 NaH_2_PO_4_, 10 Dextrose). Custom made chambers were designed and built to measure short-circuit current in 0.33 cm^2^ transwell inserts (www.thiagarajahlab/tools). The cells were maintained at 37°C, and short-circuit current was measured using an VCCMC8 multichannel voltage clamp (Physiologic Instruments), and LabChart (ADInstruments) was used to record measurements.

### Na^+^/H^+^ exchanger measurement

Na^+^/H^+^ exchanger measurements were done using a variation of previously optimized protocols (29, 47). Briefly, enteroids were plated onto coverslips and cultured under differentiation media conditions (Supplementary Methods). Cells were loaded with SNARF-5F-AM (5 μM) in 50 mM NH_4_Cl-prepulse buffer for 15 min at 37 °C, 5% CO2. Coverslips were mounted in a preheated chamber (37 °C, Warner Instruments) and imaging was done using a 20X air objective on a Zeiss 880 laser scanning microscope at 488 nm excitation, 580/640nm emission. For each condition, fluid was aspirated and replaced with 1 mL fresh buffer maintained at 37 °C. Images were collected after incubation in Na^+^-free tetramethylammonium chloride (TMA) buffer and at various intervals in modified Krebs Na^+^ buffer. Intracellular pH was calibrated at the end of each experiment by exposure to 20 μM nigericin for 10 min in K+ clamp buffers to set the pH at 6, 7, and 8.

### Gene expression analysis by qPCR

Total RNA was extracted from cells using the RNeasy Mini Kit (Qiagen). Cell pellets were lysed in Buffer RLT and processed according to the manufacturer′s protocol. Total RNA concentrations were measured by absorbance at 260 nm, and quality was assessed by A260/A280 ratios. cDNA was synthesized from 1µg of RNA, including DNA elimination step, using QuantiTect Reverse Transcription Kit (Qiagen) according to manufacturer’s protocol.

Target transcripts were amplified using the primers listed below (Integrated DNA Technologies, Inc.) and Sso Advanced Universal SYBR Green Supermix according to the manufacturer′s protocol (BioRad). All qPCR reactions were assayed in triplicate for each sample, and the average Cq value was used to calculate the mean expression ratio of the test sample compared with the control sample using the 2-ΔΔCt method. Cq values for targets were analyzed relative to Cq values for the GAPDH housekeeping gene. The sequences for the qPCR primers were designed using Primerbank (Supplementary Methods) and purchased from Integrated DNA technologies (IDT, Iowa USA).

### Bulk RNA sequencing Analysis

The paired RNA-seq reads were preprocessed using FASTP. The filtered reads were then mapped to the GRCh41 genome assembly using HISAT2 with an overall alignment greater than 90%. The mapped transcripts were quantified using Salmon and differential analysis was performed using DESEQ2 with a significance FDR cut-off of 0.05. All subsequent analysis and filtering were carried out using custom written programs in Matlab (48–51).

### Statistics

Significance was assessed using two-tailed t-test or two-way ANOVA with post-hoc multiple comparison testing (Tukey-Kramer) and where indicated p<0.05 was considered significant. Graphs were generated using GraphPad Prism 8.

## ACKNOWLEDGEMENTS

The authors thank the patients and their families who consented and participated in this study, as well as all healthcare providers, particularly members of the Boston Children’s Congenital Enteropathy and Home Parenteral Nutrition Programs. We would also like to thank Mr Kyle Smith in the HDDC Imaging Core for assistance with electron microscopy imaging.

## AUTHOR CONTRIBUTIONS

Conceptualization – J.R.T., J.R.G, I.K.; methodology – J.R.T., K.R., H.O., J.T.R, M.T., S.H.; clinical investigation and data – L.J., M.S., J.D.G.; investigation – M.K., K.R., H.O., M.T., E.K., I.K., J.R.T.; writing – original draft, J.R.T., J.R.G., I.K.; writing – review & editing, all authors; supervision – J.R.G., I.K., S.H., J.R.T.

---

**SUPPLEMENTARY FIGURE 1:**
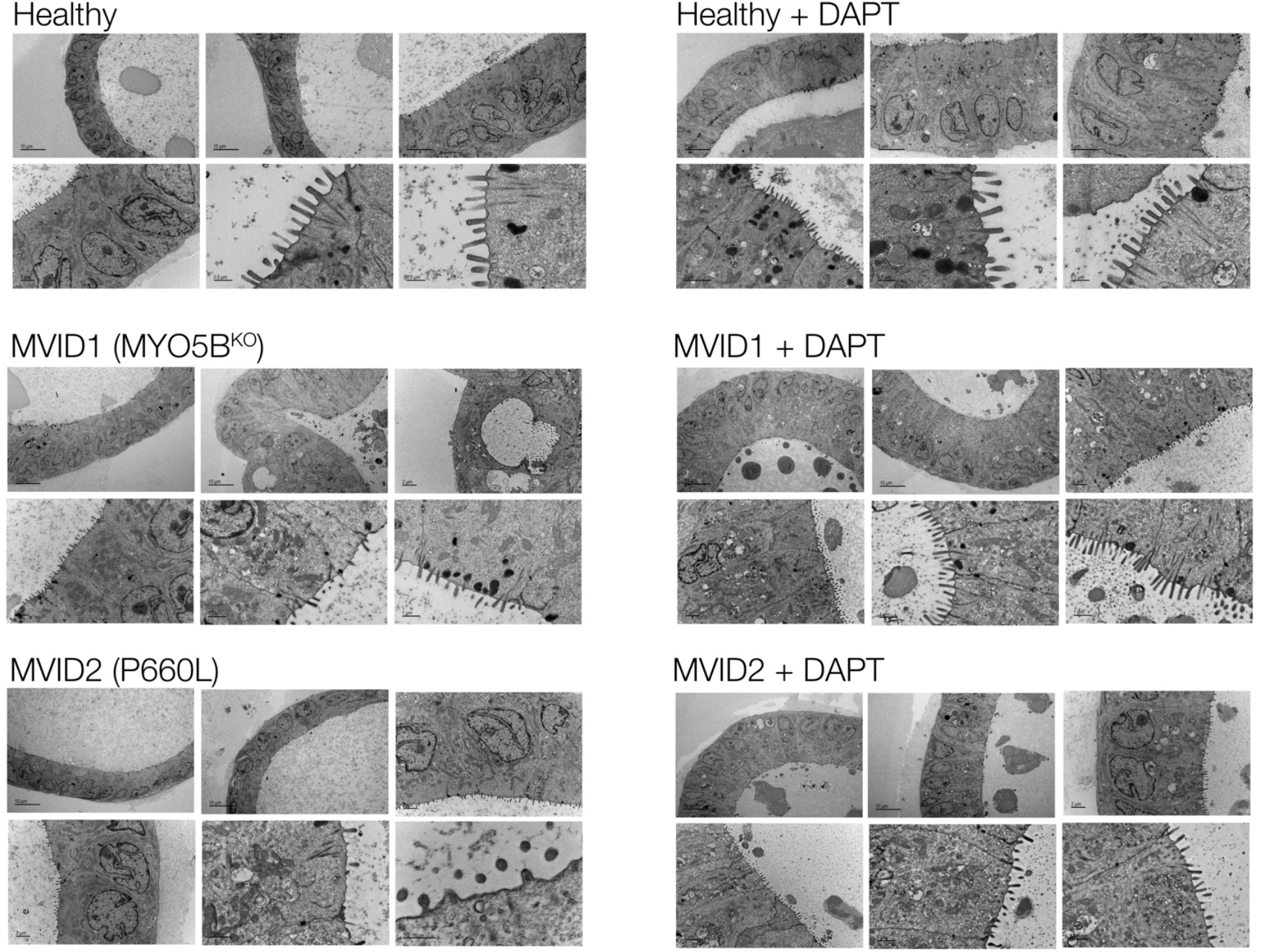
Representative electron micrographs of healthy and MVID enteroids ± DAPT (10 μM).

**SUPPLEMENTARY FIGURE 2:**
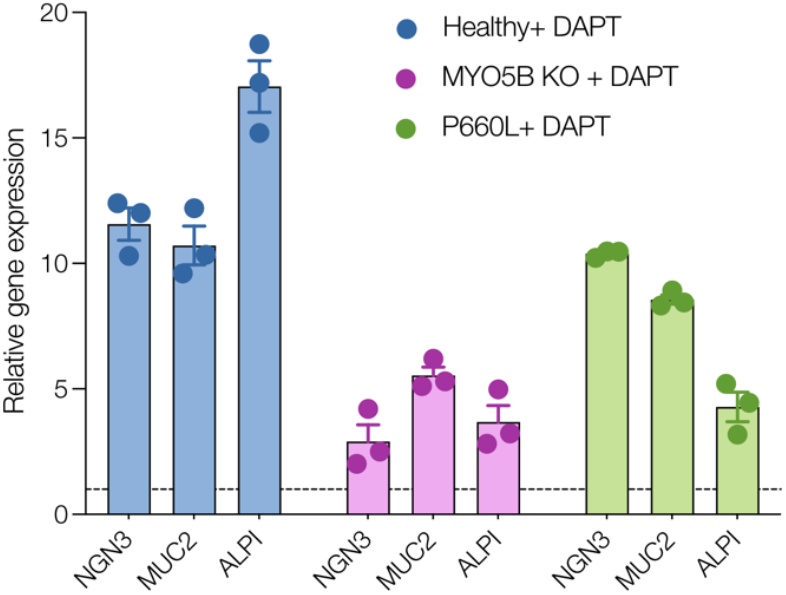
Relative gene expression (normalized to differentiated) for neurogenin3 (NGN), mucin 2 (MUC2) and alkaline phosphatase (ALPI) in healthy and MVID enteroids following addition of DAPT. Dotted line indicates baseline without DAPT. Error bars represent means ± SEM, n=3 experiments.

**SUPPLEMENTARY FIGURE 3:**
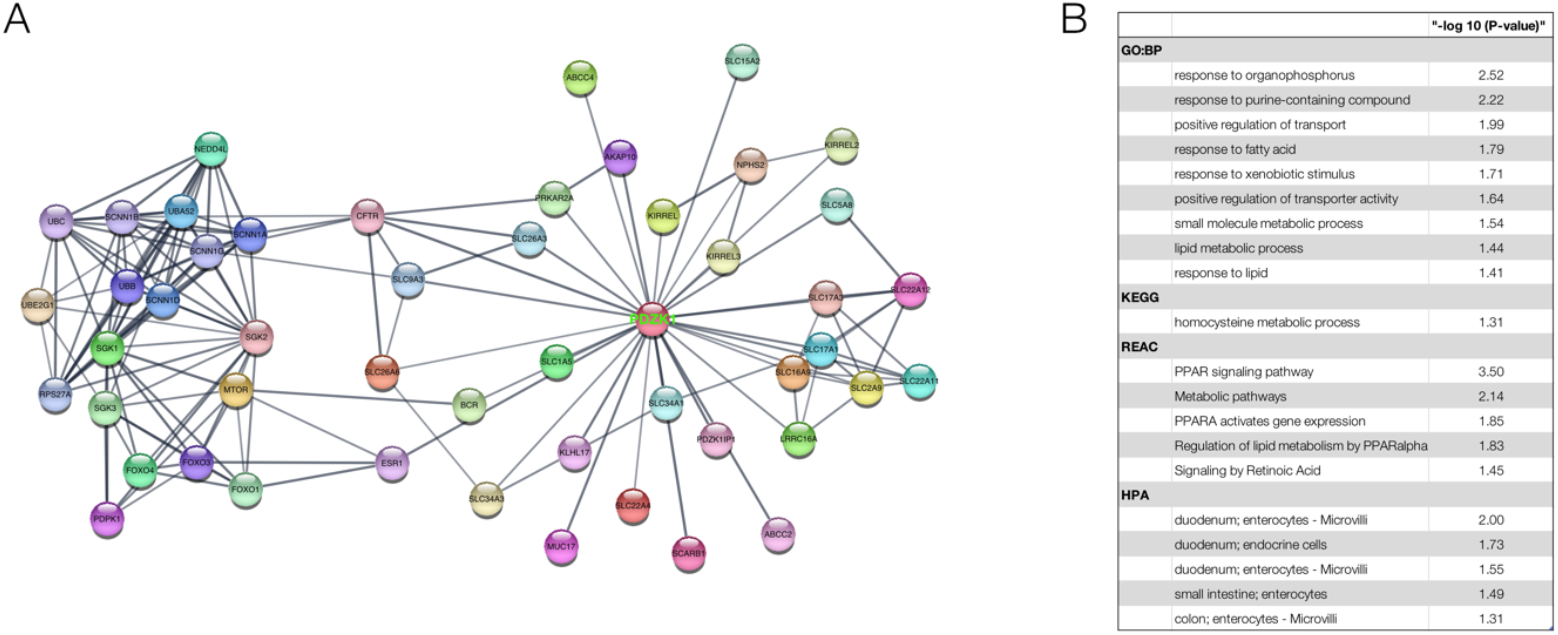
**A**. STRING protein interaction network analysis for SGK2 and PDZK1. **B**.Pathway analysis showing most significant GO terms, HPA terms and KEGG pathways with P-values.

**SUPPLEMENTARY MOVIE 1:**
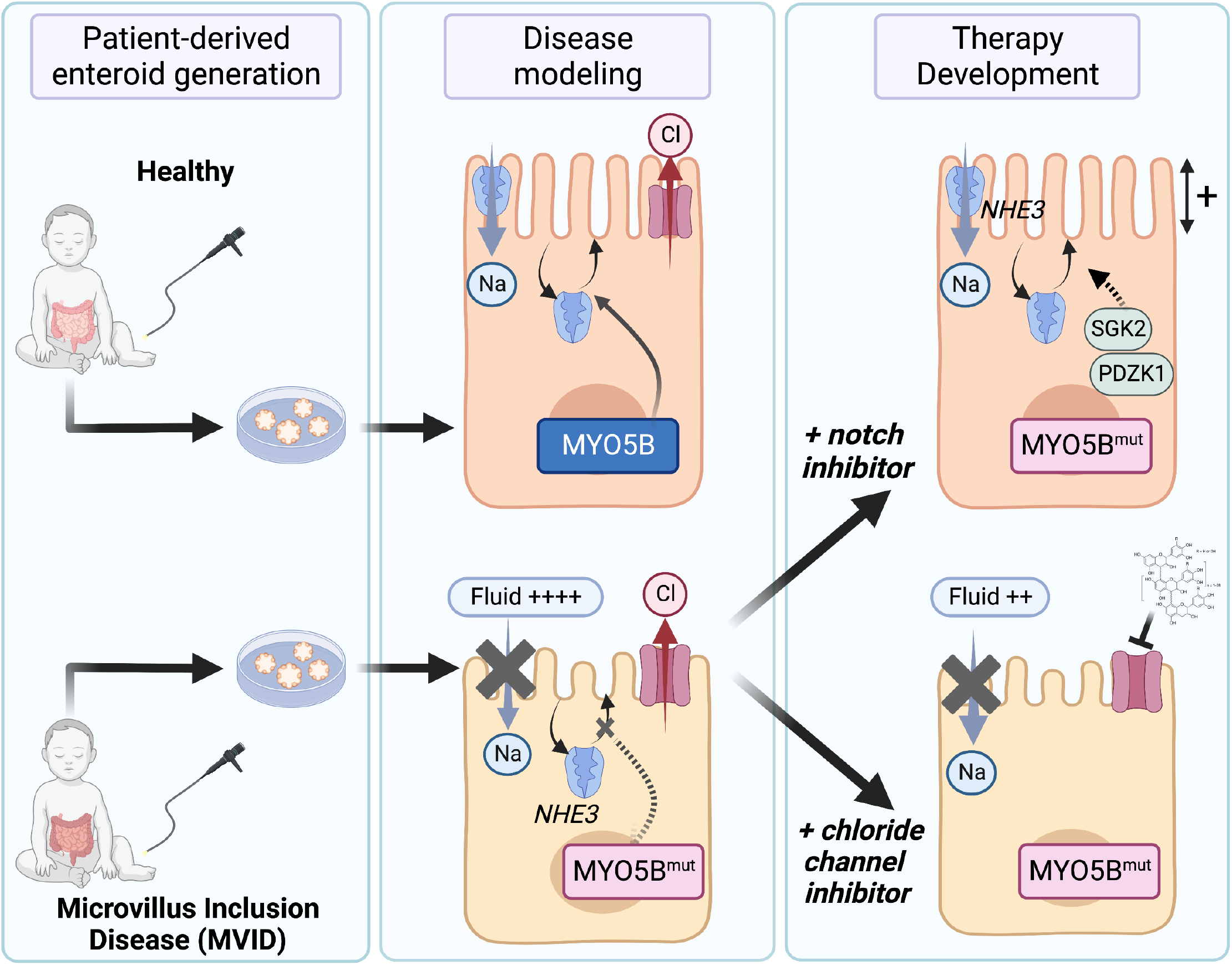
Lightsheet scan showing three-dimensional MVID enteroid stained for actin (white) and nuclei (blue) indicating abnormal intracellular large inclusions with microvilli.

## REFERENCES

1. Müller T et al. MYO5B mutations cause microvillus inclusion disease and disrupt epithelial cell polarity. Nat Genet 2008;40(10):1163–1165.

2. van der Velde KJ et al. An overview and online registry of microvillus inclusion disease patients and their MYO5B mutations. Hum Mutat 2013;34(12):1597–1605.

3. Bowman DM, Kaji I, Goldenring JR. Altered MYO5B Function Underlies Microvillus Inclusion Disease: Opportunities for Intervention at a Cellular Level. Cell Mol Gastroenterol Hepatol 2022;14(3):553–565.

4. Babcock SJ, Flores-Marin D, Thiagarajah JR. The genetics of monogenic intestinal epithelial disorders [Internet]. Hum Genet [published online ahead of print: November 23, 2022]; doi:10.1007/s00439-022-02501-5

5. Erickson RP, Larson-Thomé K, Valenzuela RK, Whitaker SE, Shub MD. Navajo microvillous inclusion disease is due to a mutation in MYO5B. Am. J. Med. Genet. A 2008;146A(24):3117–3119.

6. Ruemmele FM, Schmitz J, Goulet O. Microvillous inclusion disease (microvillous atrophy). Orphanet J Rare Dis 2006;1:22.

7. Halac U et al. Microvillous inclusion disease: how to improve the prognosis of a severe congenital enterocyte disorder. J Pediatr Gastroenterol Nutr 2011;52(4):460–465.

8. Jayawardena D, Alrefai WA, Dudeja PK, Gill RK. Recent advances in understanding and managing malabsorption: focus on microvillus inclusion disease. F1000Res 2019;8:F1000 Faculty Rev-2061.

9. Lapierre LA, Goldenring JR. Interactions of myosin vb with rab11 family members and cargoes traversing the plasma membrane recycling system. Methods Enzymol 2005;403:715–723.

10. Knowles BC et al. Myosin Vb uncoupling from RAB8A and RAB11A elicits microvillus inclusion disease. J Clin Invest 2014;124(7):2947–2962.

11. Roland JT et al. Rab GTPase-Myo5B complexes control membrane recycling and epithelial polarization. Proc Natl Acad Sci U S A 2011;108(7):2789–2794.

12. Vogel GF et al. Cargo-selective apical exocytosis in epithelial cells is conducted by Myo5B, Slp4a, Vamp7, and Syntaxin 3. J Cell Biol 2015;211(3):587–604.

13. Schneeberger K et al. An inducible mouse model for microvillus inclusion disease reveals a role for myosin Vb in apical and basolateral trafficking. Proc Natl Acad Sci U S A 2015;112(40):12408–12413.

14. Lapierre LA et al. Myosin vb is associated with plasma membrane recycling systems. Mol Biol Cell 2001;12(6):1843–1857.

15. Engevik AC et al. Loss of MYO5B Leads to Reductions in Na+ Absorption With Maintenance of CFTR-Dependent Cl-Secretion in Enterocytes. Gastroenterology 2018;155(6):1883-1897.e10.

16. Kaji I et al. Lysophosphatidic Acid Increases Maturation of Brush Borders and SGLT1 Activity in MYO5B-deficient Mice, a Model of Microvillus Inclusion Disease. Gastroenterology 2020;159(4):1390-1405.e20.

17. Kravtsov D et al. Myosin 5b loss of function leads to defects in polarized signaling: implication for microvillus inclusion disease pathogenesis and treatment. Am. J. Physiol. Gastrointest. Liver Physiol. 2014;307(10):G992–G1001.

18. Engevik AC et al. Editing Myosin VB Gene to Create Porcine Model of Microvillus Inclusion Disease, With Microvillus-Lined Inclusions and Alterations in Sodium Transporters. Gastroenterology 2020;158(8):2236-2249.e9.

19. Sato T, Clevers H. Growing self-organizing mini-guts from a single intestinal stem cell: mechanism and applications. Science 2013;340(6137):1190–1194.

20. Thiagarajah JR et al. Advances in Evaluation of Chronic Diarrhea in Infants. Gastroenterology [published online ahead of print: April 12, 2018]; doi:10.1053/j.gastro.2018.03.067

21. Dekkers JF et al. A functional CFTR assay using primary cystic fibrosis intestinal organoids. Nat Med 2013;19(7):939–945.

22. Jardine S et al. Drug Screen Identifies Leflunomide for Treatment of Inflammatory Bowel Disease Caused by TTC7A Deficiency. Gastroenterology 2020;158(4):1000–1015.

23. Beumer J, Clevers H. Cell fate specification and differentiation in the adult mammalian intestine. Nat Rev Mol Cell Biol 2021;22(1):39–53.

24. Kaji I et al. Cell differentiation is disrupted by MYO5B loss through Wnt/Notch imbalance. JCI Insight 2021;6(16):150416.

25. Sato T et al. Long-term expansion of epithelial organoids from human colon, adenoma, adenocarcinoma, and Barrett’s epithelium. Gastroenterology 2011;141(5):1762–1772.

26. VanDussen KL et al. Development of an enhanced human gastrointestinal epithelial culture system to facilitate patient-based assays. Gut 2015;64(6):911–920.

27. Burman A et al. Modeling of a novel patient-based MYO5B point mutation reveals insights into MVID pathogenesis. Cell Mol Gastroenterol Hepatol 2022;S2352-345X(22)00266–1.

28. Thiagarajah JR, Broadbent T, Hsieh E, Verkman AS. Prevention of toxin-induced intestinal ion and fluid secretion by a small-molecule CFTR inhibitor. Gastroenterology 2004;126(2):511–519.

29. Foulke-Abel J et al. Human Enteroids as a Model of Upper Small Intestinal Ion Transport Physiology and Pathophysiology. Gastroenterology 2016;150(3):638-649.e8.

30. Tradtrantip L, Namkung W, Verkman AS. Crofelemer, an antisecretory antidiarrheal proanthocyanidin oligomer extracted from Croton lechleri, targets two distinct intestinal chloride channels. Mol. Pharmacol. 2010;77(1):69–78.

31. Kobayashi T, Deak M, Morrice N, Cohen P. Characterization of the structure and regulation of two novel isoforms of serum- and glucocorticoid-induced protein kinase. Biochem J 1999;344 Pt 1(Pt 1):189–197.

32. Lang F, Stournaras C, Zacharopoulou N, Voelkl J, Alesutan I. Serum- and glucocorticoid-inducible kinase 1 and the response to cell stress. Cell Stress 2018;3(1):1–8.

33. Pao AC. SGK regulation of renal sodium transport. Curr Opin Nephrol Hypertens 2012;21(5):534–540.

34. Pao AC et al. Expression and role of serum and glucocorticoid-regulated kinase 2 in the regulation of Na+/H+ exchanger 3 in the mammalian kidney. Am J Physiol Renal Physiol 2010;299(6):F1496–1506.

35. Maiyar AC, Huang AJ, Phu PT, Cha HH, Firestone GL. p53 Stimulates Promoter Activity of the sgk Serum/Glucocorticoid-inducible Serine/Threonine Protein Kinase Gene in Rodent Mammary Epithelial Cells (∗). Journal of Biological Chemistry 1996;271(21):12414–12422.

36. Imaizumi K, Tsuda M, Wanaka A, Tohyama M, Takagi T. Differential expression of sgk mRNA, a member of the Ser/Thr protein kinase gene family, in rat brain after CNS injury. Molecular Brain Research 1994;26(1):189–196.

37. Webster MK, Goya L, Firestone GL. Immediate-early transcriptional regulation and rapid mRNA turnover of a putative serine/threonine protein kinase. Journal of Biological Chemistry 1993;268(16):11482–11485.

38. Sangphech N, Palaga T. Cross-regulation of notch/AKT and serum/glucocorticoid regulated kinase 1 (SGK1) in IL-4-stimulated human macrophages. Int Immunopharmacol 2021;101(Pt A):108312.

39. Xu D, Huang H, Toh MF, You G. Serum- and glucocorticoid-inducible kinase sgk2 stimulates the transport activity of human organic anion transporters 1 by enhancing the stability of the transporter. Int J Biochem Mol Biol 2016;7(1):19–26.

40. Chun J et al. The Na(+)/H(+) exchanger regulatory factor 2 mediates phosphorylation of serum- and glucocorticoid-induced protein kinase 1 by 3-phosphoinositide-dependent protein kinase 1. Biochem Biophys Res Commun 2002;298(2):207–215.

41. Bultema JJ, Di Pietro SM. Cell type-specific Rab32 and Rab38 cooperate with the ubiquitous lysosome biogenesis machinery to synthesize specialized lysosome-related organelles. Small GTPases 2013;4(1):16–21.

42. Ohbayashi N, Fukuda M, Kanaho Y. Rab32 subfamily small GTPases: pleiotropic Rabs in endosomal trafficking. J Biochem 2017;162(2):65–71.

43. Waschbüsch D et al. Rab32 interacts with SNX6 and affects retromer-dependent Golgi trafficking. PLoS One 2019;14(1):e0208889.

44. Lamprecht G, Seidler U. The emerging role of PDZ adapter proteins for regulation of intestinal ion transport. American Journal of Physiology-Gastrointestinal and Liver Physiology 2006;291(5):G766–G777.

45. LaLonde DP, Garbett D, Bretscher A. A regulated complex of the scaffolding proteins PDZK1 and EBP50 with ezrin contribute to microvillar organization. Mol Biol Cell 2010;21(9):1519–1529.

46. Avula LR, Chen T, Kovbasnjuk O, Donowitz M. Both NHERF3 and NHERF2 are necessary for multiple aspects of acute regulation of NHE3 by elevated Ca2+, cGMP, and lysophosphatidic acid. Am J Physiol Gastrointest Liver Physiol 2018;314(1):G81–G90.

47. Levine SA, Montrose MH, Tse CM, Donowitz M. Kinetics and regulation of three cloned mammalian Na+/H+ exchangers stably expressed in a fibroblast cell line. J Biol Chem 1993;268(34):25527–25535.

48. Chen S, Zhou Y, Chen Y, Gu J. fastp: an ultra-fast all-in-one FASTQ preprocessor. Bioinformatics 2018;34(17):i884–i890.

49. Patro R, Duggal G, Love MI, Irizarry RA, Kingsford C. Salmon provides fast and bias-aware quantification of transcript expression. Nat Methods 2017;14(4):417–419.

50. Love MI, Huber W, Anders S. Moderated estimation of fold change and dispersion for RNA-seq data with DESeq2. Genome Biol 2014;15(12):550.

51. Kim D, Paggi JM, Park C, Bennett C, Salzberg SL. Graph-based genome alignment and genotyping with HISAT2 and HISAT-genotype. Nat Biotechnol 2019;37(8):907–915.

