## Supplemental Methods for "Therapy Development for Microvillus Inclusion Disease using Patient-derived Enteroids"

### Supplementary Materials and Methods

#### Patient Clinical Summaries

**Patient 1:** Male with microvillus inclusion disease who is sustained on parenteral nutrition (PN). Patient was diagnosed within 2 months of age secondary to intractable diarrhea with need for PN support. Endoscopic evaluation was notable for villus blunting and abnormal CD10 brush border staining. Patient's medical course has been complicated by oral aversion, central line associated bloodstream infection, iron deficiency anemia, electrolyte derangements, and PN associated liver disease. Endoscopic biopsies for enteroid generation obtained at age 3 yrs. His PN support requires sodium supplementation (ranging from 14-17 mEq/kg/day) as well as maximum acetate provision and potassium infusion (0.25 mEq/kg/h rate due to high gastrointestinal losses.

**Patient 2:** Female born at 39 weeks gestation with onset of severe secretory diarrhea within the 1st week of life associated with acidosis and severe dehydration. EGD performed at 1 month consistent with MVID (gross blunting of villi, microvillous inclusions on EM). Patient remained PN dependent until bowel transplant (age 5) with biopsies retained for enteroid generation. Underwent multivisceral transplant (liver, pancreas, small intestine and colon) with retained native duodenum. Patient alive 4 years post-transplant.

#### Enteroid Media and Culture

*Expansion Media:*

| Component | Volume | Catalog # | Final Concentration |
|---|---|---|---|
| L-WRN Conditioned Media | 65 mL | ATCC CRL-3276 | 65% |
| Advanced DMEM/F12 | 30 mL | Gibco 12634-028 | 30% |
| GlutaMax (100X) | 1 mL | Gibco 35050-061 | 1% |
| HEPES 1M | 1 mL | Gibco 15630-080 | 10 mM |
| Primocin | 200 µL | Invivogen ant-pm-2 | 0.2% |
| Normocin | 200 µL | Invivogen ant-nr-2 | 0.2% |
| B27 | 1 mL | Gibco 12587010 | 1% |
| N2 | 500 µL | Gibco 17502-048 | 0.5% |
| Nicotinamide 1M | 1 mL | Sigma N0636 | 10 mM |

| | | | |
|---|---|---|---|
| N-Acetyl-Cystein (500 mM) | 100 µL | Sigma A8199 | 500 µM |
| A 83-01 (500 µM) | 100 µL | Sigma SML0788 | 500 nM |
| SB202190 (5 mg/505 µL) | 33.2 µL | Sigma S7067 | |
| EGF (500 µg/mL) | 10 µL | Peprotech 315-09 | 50 ng/mL |
| Gastrin (500 µM) | 10 µL | Sigma G9145 | 10 nM |
| Prostaglandin E2 (5 mg/mL) | 1 µL | Sigma P5640 | 100 nM |
| Y-27632 (3.2 mg/mL) | 100 µL | Sigma Y0503 | 10 µM |
| **Total Volume** | **100 mL** | | |

*Differentiation Media:*

| **Component** | **Volume** | **Catalog #** | **Final Concentration** |
|---|---|---|---|
| L-WRN Conditioned Media | 15 mL | ATCC CRL-3276 | 15% |
| Advanced DMEM/F12 | 80 mL | Gibco 12634-028 | 80% |
| GlutaMax (100X) | 1 mL | Gibco 35050-061 | 1% |
| HEPES 1M | 1 mL | Gibco 15630-080 | 10 mM |
| Primocin | 200 µL | Invivogen ant-pm-2 | 0.2% |
| Normocin | 200 µL | Invivogen ant-nr-2 | 0.2% |
| B27 | 1 mL | Gibco 12587010 | 1% |
| N2 | 500 µL | Gibco 17502-048 | 0.5% |
| Nicotinamide 1M | 1 mL | Sigma N0636 | 10 mM |
| N-Acetyl-Cystein (500 mM) | 100 µL | Sigma A8199 | 500 µM |
| EGF (500 µg/mL) | 10 µL | Peprotech 315-09 | 50 ng/mL |
| **Total Volume** | **100mL** | | |

*Plating on Transwells*:

Formed enteroids were removed from tissue culture plates and Matrigel® was dissolved in Cell-recovery solution (Corning). Enteroids were dissociated by vigorous pipetting and incubation at 37ºC with TRIPL-E (ThermoFisher) for 2-3 mins. Cells were plated onto human placental collagen

IV [Please confirm collagen type] (Sigma)-coated Transwell filters (Corning) with 0.3-µm-pore size inserts and cultured for 2-4 days in Expansion media including Rho Kinase inhbitor (Y-27632) until transepithelial resistance (TEER) started to rise. Media was switched to the differentiation media and electrophysiological and immunohistological measurements done after 10-12 days, when TEER reached to >2000Ω/cm$^2$.

*Plating on Coverslips:*

Formed enteroids were removed from tissue culture plates and Matrigel® was removed in Cell-recovery solution. Enteroids were dissociated by vigorous pipetting and incubation at 37ºC with TRIPL-E for 2-3 mins. Cells were plated onto human placental collagen coated coverslips and cultured in the differentiation media ± DAPT for four days.

**Enteroid Formation Assay**

Formed enteroids (P1-3) were removed from tissue culture plates and Matrigel® was removed in cell-recovery solution. Enteroids were dissociated by vigorous pipetting and incubation at 37ºC with TRIPL-E for 2-3mins. Cells were counted and re-plated in Matrigel® for MVID and healthy enteroids at approximately same density. Cells were cultured in the expansion media for 3 days and formed enteroid numbers were counted in each plate well.

**Enteroid Swelling Assay**

Enteroid swelling after Crofelemer (100 µM) or vehicle treatment (PBS) was performed as previously described (1). Measurements of cell diameter was facilitated by Image J.

**EM image analysis**

EM images (at least 10-15 per enteroid) were analyzed blinded in Image J for measurement of microvilli and actin bundle length and distance between apical membrane and the majority of cell organelles.

**qPCR primer sequences**

<u>PCR primer sequences:</u>

Human SGK2;

Primer 1: 5'- CCACGGACTTCGACTTCCTC -3'

Primer 2: 5'- GTGCCGCACGTTCTTCAGA -3'

Human PDZK1:

PrimeTime™ Predesigned qPCR Assays (Assay Id: Hs.PT.58.4953162)

Human RAB32:

Primer 1: 5'- CAGGTGGACCAATTCTGCAAA -3'

Primer 2: 5'- GGCAGCTTCCTCTATGTTTATGT -3'

**MxIF Antibodies**

| | | | |
|---|---|---|---|
| ACTG1 | sc-65638 AF488 | AB_2890619 | 1:100 |
| Beta-catenin | NBP1-54467IR | 12F7 | 1:50 |
| CD10 | sc-46656 AF488 | AB_2890648 | 1:100 |
| CHGA | NBP2-47850IR | CGA/493 | 1:2000 |
| Defensin 5A | NB110-60002IR | 8c8 | 1:200 |
| Cd26/DPP4 | NBP2-70588C | OTI11D7 | 1:200 |
| EGFR pY1068 | ab205828 | AB_2890267 | 1:200 |
| Ep-CAM | ab275122 | EPR677(2) | 1:100 |
| GLUT2 | NBP2-22218AF647 | AB_2890913 | 1:50 |
| LAMP2A | ab282009 | EPR4207(2) | 1:50 |
| MYO5B | NBP1-87746 | AB_11034537 | 5 $\mu$g/ml (Zenon labeled) |
| pNHE3 | NB110-81529R | 14D5 | 1:50 |
| SGLT1 | NBP2-38748 | AB_2890609 | 5 $\mu$g/ml (Zenon labeled) |
| Villin | sc-58897 AF488 | 1D2C3 | 1:50 |

**SUPPLEMENTARY FIGURE 1:** Representative electron micrographs of healthy and MVID enteroids ± DAPT (10 $\mu$M).

**SUPPLEMENTARY FIGURE 2:** Relative gene expression (normalized to differentiated) for neurogenin3 (NGN), mucin 2 (MUC2) and alkaline phosphatase (ALPI) in healthy and MVID enteroids following addition of DAPT. Dotted line indicates baseline without DAPT. Error bars represent means ± SEM, n=3 experiments.

**SUPPLEMENTARY FIGURE 3:** **A.** STRING protein interaction network analysis for SGK2 and PDZK1. **B.** Pathway analysis showing most significant GO terms, HPA terms and KEGG pathways with P-values.

**SUPPLEMENTARY MOVIE 1:** Lightsheet scan showing three-dimensional MVID enteroid stained for actin (white) and nuclei (blue) indicating abnormal intracellular large inclusions with microvilli.
